# Pangenomic Landscapes Shape Performances of a Synthetic Genetic Circuit Across *Stutzerimonas* Species

**DOI:** 10.1101/2024.02.15.580380

**Authors:** Dennis Tin Chat Chan, Hans C. Bernstein

## Abstract

Engineering identical genetic circuits into different species typically results in large differences in performance due to the unique cellular environmental context of each host, a phenomenon known as the “chassis-effect”. A better understanding of how genomic and physiological contexts underpin the chassis-effect will greatly improve biodesign strategies across diverse microorganisms. Here, we combined a pangenomics-based gene expression analysis with quantitative measurements of performance from an engineered genetic inverter device to uncover how genome structure and function relates to the observed chassis-effect across six closely related *Stutzerimonas* hosts. Our results reveal that genome architecture underpins divergent responses between our chosen non-model bacterial hosts to engineered genetic circuits. Specifically, differential expression of the core genome, gene clusters shared between all hosts, were found to be the main source of significant concordance to the observed genetic device performance, whereas specialty genes from respective accessory genomes were not significant. A data-driven investigation revealed that genes involved in denitrification and components of trans-membrane transporter proteins were among the most differentially expressed gene clusters in response to the genetic device. Our results show the chassis-effect can be traced along differences among genome-encoded functions that are mostly conserved and that these differences create a unique biodesign space among closely related species.

**GRAPHICAL ABSTRACT:** 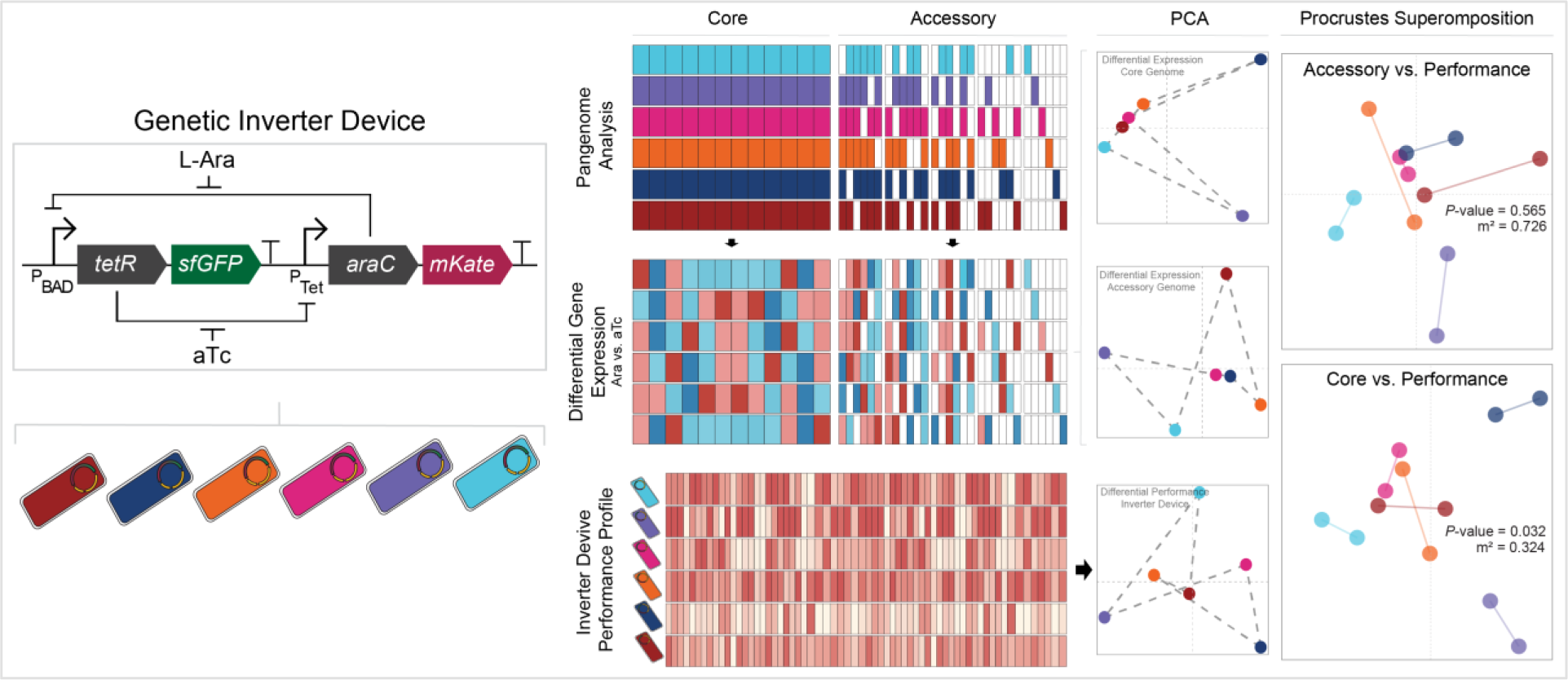

## IMPORTANCE

Contemporary synthetic biology endeavors often default to a handful of model organisms to host their engineered systems. Model organisms such as *Escherichia coli* serve as attractive hosts due to their tractability but do not necessarily provide the ideal environment to optimize performance, and as more novel microbes are domesticated for use at biotechnology platforms, synthetic biologists are urged to explore the chassis-design space in order to optimize their systems and deliver on the promises of synthetic biology. Therefore, the consequences of the chassis-effect will only become more relevant as the field of biodesign grows. In our work, we demonstrate that the performance of a genetic device is highly dependent on the host environment it operates within, promoting the notion that the chassis can be considered a design variable to tune circuit function. Importantly, our results unveil that the chassis-effect can be traced along similarities in genome architecture, specifically the shared core genome. Our study advocates for the exploration of the chassis-design space and is a step forward to empowering synthetic biologists with knowledge for more efficient exploration of the chassis-design space to enable the next generation of broad-host-range synthetic biology.

## INTRODUCTION

As synthetic biologists continue to explore the design space of engineered genetic circuits, we are presented with a complex landscape where functional fidelity depends on host physiology, environment, and genetic tractability^1,2^. This has given rise to a subdiscipline of broad-host-range synthetic biology, which aims to create versatile genetic systems that can operate across diverse organisms, and is particularly beneficial towards biotechnologies that capitalize on microbial diversity^3,4^. Despite the plethora of modular genetic parts available, we are still faced with the challenge that identical genetic devices exhibit consequentially different performance depending on the host context the device operates in, a process termed the “chassis-effect”^5,6^ or “context dependence”^7,8^. The chassis-effect hinders the predictability of circuit function based on part composition alone and can cause any circuit optimization typically done in a model organism (e.g., *Escherichia coli*) to be rendered null once introduced into a different host environment^9^. This added layer of instability limits the current state of microbial biodesign and biases our understanding towards model organisms, even though more suitable hosts may exist for a given application^10–12^. Overall, the chassis-effect constrains the design-build-test cycle by demanding costly repetitions of trial-and-error experimentation. There remain major knowledge gaps as to which cellular processes underpin species-specific chassis-effects. Closing this knowledge gap undoubtedly enables more efficient engineering of novel hosts and consequently shifts the current engineering dogma towards the notion that the host itself can be considered a part to tune circuit functions^7,8^, which is a paradigm shift towards a new understanding of different microbial species and their unique genomes as customized hardware for biodesign^13^.

Previous efforts to address this knowledge gap has shown that device performances between bacterial hosts can be better explained by the differences in the growth physiology of hosts rather than genomic or phylogenetic relatedness^14^. In turn, the available physiological states for a given host are ultimately shaped by those functions encoded in their genomes, and perhaps more importantly, the expression of gene products that control cellular physiology. Deeper investigation into the gene expression responses towards the activity of a heterologous genetic circuits are needed to uncover insight into which cellular processes underpin observable chassis-effects. Designing such a study is difficult, as it requires a suit of microbial hosts capable of operating an identical genetic device, produce an observable chassis-effect under the same growth conditions and share significant enough amount of genomic identity for their interspecies gene expression to be comparable. We developed a synthetic biology kit for the *Stutzerimonas* genus to specifically address this challenge. Recent re-examination of the *Pseudomonas* genus invigorated with high quality genome sequencing data led to the delineation of the clade into several novel genus^15,16^, one such genus being *Stutzerimonas*^17,18^. Several members of the *Stutzerimonas* with sequenced genomes are available in culture collections and previous studies has highlighted the natural high transformation competence of *Stutzerimonas* spp^19,20^. Furthermore, many *Stutzerimonas* members have innate phenotypes that have garnered attention as potential microbial cell factories and/or bioremediation agents^21–24^, making them appealing targets for domestication and biodesign applications.

In this work, we transformed an inducible genetic inverter device into six *Stutzerimonas* hosts that share sufficient amount of genomic identify to form a quality pangenome^17^, consisting of a core (i.e., orthologous genes shared between all hosts) and an accessory genome (i.e., orthologous genes shared only by a subset of hosts or genes unique to host). We comparatively quantified an observable chassis-effect in device performance within these closely related *Stutzerimonas* hosts and sequenced the global transcriptomes during different operation modes of the inverter. This investigative workflow enabled us to ask the guiding scientific question of whether the observed chassis-effect is more influenced by expression of conserved core genes that are fundamental to growth physiology and cellular housekeeping or by less conserved functions encoded within the accessory genome of our experimental platform. We were also able to ask if unique, species-specific gene expression patterns can serve as a concordant predictor of device performance. Our results show significant correlation between transcriptional response of shared core genes and inverter performance, as well as between growth physiology and inverter performance, suggesting the mechanism in which core genetic elements contribute to the chassis-effect effect is through changes in basic cellular physiology related to growth.

## RESULTS

### A *Stutzerimonas* tool kit for pangenome-guided synthetic biology

Connecting structures and functions of multiple genomes to a measurable chassis-effect requires a standardized, broad-host-range genetic circuit. We therefore implemented a modified build from one of our previously described genetic inverters^14^ (Fig. 1a). This inverter can be directionally induced (toggled) by anhydrotetracycline (aTc) and L-arabinose (Ara) and was cloned into plasmid pS5. The circuit features two inducible promoters, pBAD and pTet, with each promoter regulating the expression of the other’s cognate transcription factor as well as a fluorescent reporter protein in a bicistronic manner. Fifteen *Stutzerimonas* hosts were screened for plasmid transformability and inverter operability, in which ten were successfully transformed via electroporation with a pBBR1-KanR backbone vector (BB23). Six of these ten were chosen for further study, which are as follows: *Stutzerimonas chloritidismutans* NCTC10475 (*S. chloritidismutans*)^25^, *Stutzerimonas perfectomarina* CCUG 44592 (*S. perfectomarina*)^18^, *Stutzerimonas degradans* FDAARGOS 876 (*S. degradans*)^18^, *Stutzerimonas stutzeri* 24a13 (*S. stutzeri* 24a13)^26^, *Stutzerimonas stutzeri* 24a75 (*S. stutzeri* 24a75)^26^ and *Stutzerimonas stutzeri* DSM 4166 (*S. stutzeri* DSM 4166)^27,28^.

**Figure 1.**
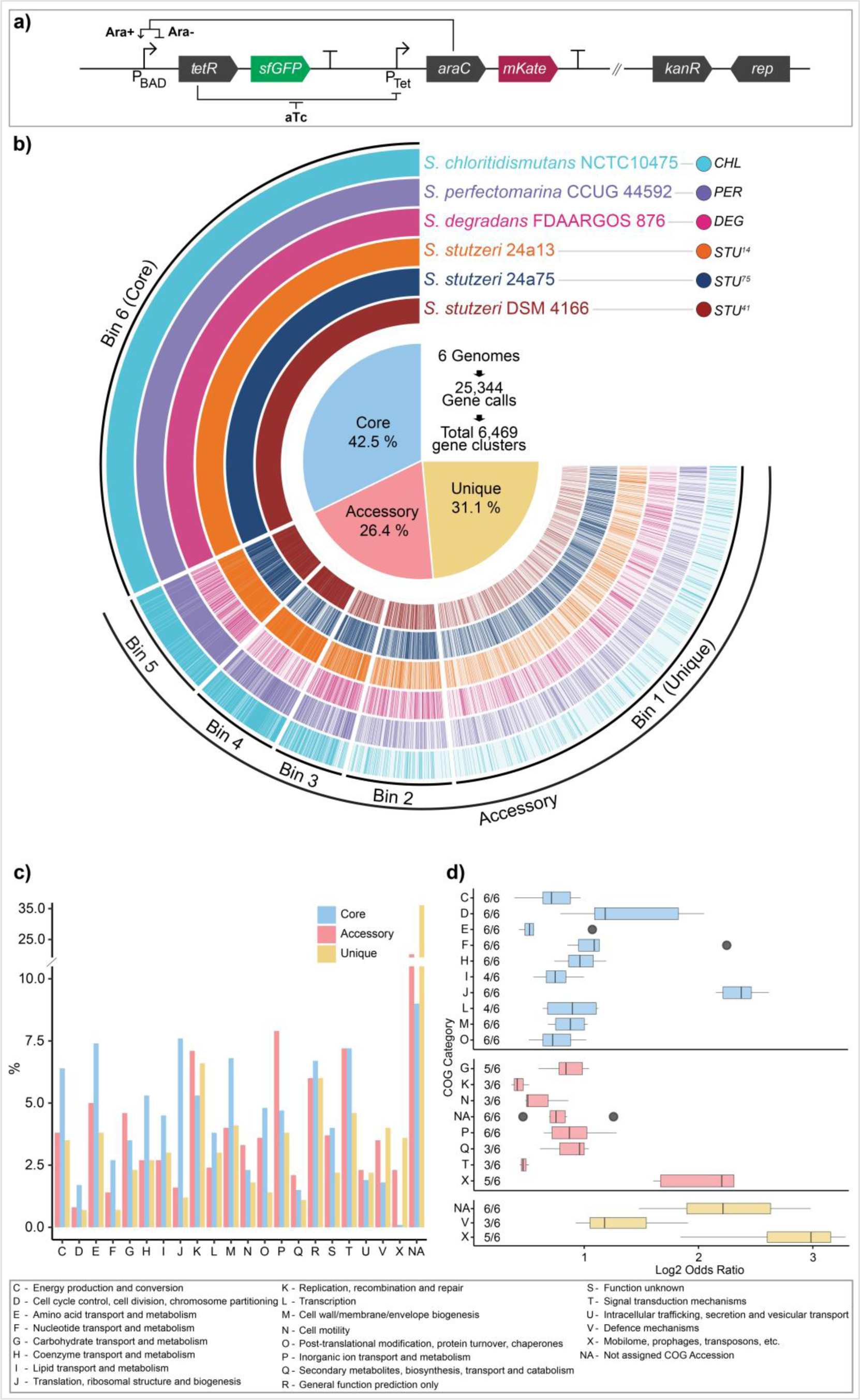
The genetic inverter and pangenome of selected *Stutzerimonas* hosts. a) Schematic representation of Ara-aTc genetic inverter composition. In presence of Ara (Ara+), Ara-bound AraC upregulates its cognate promoter (pBAD), leading to sfGFP and TetR expression and in turn creates a distinct measurable fluorescent state and leads to the downregulation of the pTet promoter. In absence of Ara (Ara-), AraC functions as a repressor. The presence of aTc leads to mKate and AraC production. The two promoters thereby act antagonistically, where upregulation of one lead to downregulation of the other. b) Gene clusters identified from pangenome analysis of the six *Stutzerimonas* hosts arranged by number of hits in host genomes where Bin 6 means all six hosts contribute with at least one gene call to the gene cluster and Bin 5 means any combination of 5 hosts contribute with at least one gene call and so on. Bins 5 to Bin 1 are further grouped together as the accessory genome, with gene clusters belonging in Bin 1 distinguished as unique. Percentages indicates portion of the 6469 gene clusters assigned to the three frequency groups (core, accessory and unique). c) Percentage COG category composition of each frequency group. d) Enrichment analysis of COG categories within each frequency group by Fisher exact test. Number “*n*/6” indicates number of hosts in which COG category was found significantly (*P*-value < 0.05, Bonferroni correction) enriched within the frequency group. Only COG categories enriched in 3 or more hosts are shown. COG category description provided at bottom of figure. CHL = *Stutzerimonas chloritidismutans* NCTC10475; DEC = *Stutzerimonas perfectomarina* CCUG 44592; DEG = *Stutzerimonas degradans* FDAARGOS 876; STU^13^ = *Stutzerimonas stutzeri* 24a13, STU^75^ = *Stutzerimonas stutzeri* 24a75; STU^41^ *Stutzerimonas stutzeri* DSM 4166.

Comparative pangenomics of the selected *Stutzerimonas* hosts revealed almost equal sizes between the core and accessory genomes, propitiously setting the stage for investigating whether the measurable chassis-effect can be attributed to differential expression from the core or accessory genomes and which specific functions covary with host-specific device performance. Pangenome analysis of *Stutzerimonas* host genomes was performed using Anvi’o^29^, leading to a total of 25,344 gene calls functionally annotated using the 2020 clusters of orthologous genes (COGs) database (Fig. 1b). Under the most sensitive settings, these gene calls were grouped into 6,469 gene clusters or “pangenomic orthologous groups”. A gene cluster is grouped into “core” or “accessory” frequency group, depending on the number of genomes the gene cluster occurs in. Core gene clusters were defined as the 42.5 % (2,751) of gene clusters that had hits across all hosts. In contrast, the “accessory” gene clusters made up the remaining 57.5 % of gene clusters that had hits in five or fewer genomes. Within the accessory genome, we further distinguish gene clusters exclusive to a single genome, referred to as “unique” gene clusters, which makes up 31.1 % (2,013) of total gene clusters. The number of host-specific gene calls ranged from 3,732 in *S. degradans* to 4,457 in *S. stutzeri* 24a13, with all hosts sharing on average 67.2 ± 4.4 % of genes, with 25.1 ± 6.3 % and 8.2 ± 3.6 % of gene clusters categorized as accessory and unique, respectively. 23.1 % of gene clusters were not assigned to any COG category (assigned to “NA” group) (Fig. 1c), with an additional 3.6 % and 4.9 % assigned to COG category S (Function Unknown) and R (General Prediction Only). 81 % of unassigned gene clusters belong in the accessory genome, with 54.7 % of these being unique gene clusters. This disproportionate number of unassigned gene clusters between the core, accessory and unique groups is consistent with previous bacterial pangenome studies^17^ and is theorized to be due to the accessory genome tending to house genes that confer a specific advantage to the organism within its niche environment and therefore might only be expressed under certain conditions, making it difficult to study such under cultivation conditions that have been standardized across multiple species.

The seemingly uneven distribution of COG categories across the three frequency groups prompted an investigation of whether certain COG categories are overrepresented (Fig. 1d). The Fisher’s exact test (*P*-value > 0.05, with Bonferroni Correction) revealed that within the core genome, COG categories consistently enriched for (i.e., significantly enriched in ≥ 3 species) are categories associated with housekeeping functions such as category C (Energy Production and Conversion), D (Cell Cycle Control, Cell Division and Chromosome Partitioning), E (Amino Acid Transport and Metabolism) and O (Post Translational Modification, Protein Turnover and Chaperones). Meanwhile, COG groups G (Carbohydrate Transport and Metabolism), P (Inorganic Ion Transport and Metabolism), Q (Secondary Metabolites Biosynthesis, Transport and Catabolism) and K (Transcription) are overrepresented in the accessory genome, suggesting the presence of metabolic pathways that confer unique nutrient assimilation capabilities. Few COG categories were found to be consistently overrepresented within the unique genome, mainly due to majority of genes being of unknown function, but the few are category X (Mobilome: Prophages and Transposons), which hints towards past viral infection events, and V (Defense Mechanisms). Other prokaryotic pangenome studies report similar pattern of enriched COG categories in their defined shared and accessory genomes^30,31^. The distinct functional identity of genomic sub-groups gives merit to the hypothesis that expression of genes among the core or accessory groups may uniquely contribute to the chassis-effect.

### The chassis-effect is observable between closely related *Stutzerimonas* hosts

Comparative measurements representing the performance of our engineered genetic inverter operated by each host under a standardized environment revealed a clear chassis-effect. Performance within *Escherichia coli* DH5α (*E*. *coli*) was also quantitatively compared as a positive control reference from a model organism. Induction response dynamics were characterized by fitting the Hill function [(*βx^n^* / (*K^n^* + *x^n^*)) + *C*] to normalized induction curves (Fig. 2a-d). Parameters *C* is the baseline output at 0 inducer concentration, *β* is the max output level at saturating input levels, *K* represents the sensitivity of the system to inducer input as well as the input responsive range, and the Hill coefficient *n* reflects the steepness of the response (varying from step-like or dosage dependent). These parameters collectively quantify interactions between inducers (aTc and Ara), their respective transcriptional factors (TetR and AraC) and the responsive operons; thereby quantitatively describing device performance. Markedly different performance profile is observed depending on the host context the genetic inverter operates from – i.e., a strong chassis-effect. For instance, the observed *K*_aTc_ values range from 1.45 to 34.24 nM aTc among the *Stutzerimonas* hosts (Fig. 2a, b), suggesting host-specific factors affecting the intracellular aTc concentration and/or different levels of TetR repressor. *E. coli* exhibited the highest sensitivity to aTc, with a *K*_aTc_ value of 0.48. The *K*_Ara_ metrics were relatively more uniform across hosts, with only an overall 3.8-fold difference between the smallest and largest *K*_Ara_ values (Fig. 2c, d). We define an additional metric, DR_T-Ara_ and DR_T-aTc_ (theoretical dynamic range) as the ratio of the estimated *β* and empirical *C* expressed as a fold-change value, which informs the theoretical possible fold-change difference in output as measured from respective reporter proteins. At saturated inducer concentrations, the highest *β*_Ara_ achieved among the *Stutzerimonas* hosts was *S. perfectomarina* at 12,500 RFU, which also achieves the highest DR_T-Ara_ value of 24.0. Overall, *E. coli* had the lowest *β*_Ara_ at 750 RFU. The range of observed *β*_aTc_ and DR_T-aTc_ values are much lower, from 2,300 RFU to 5,900 RFU, and 2.0 and 3.8, respectively.

**Figure 2.**
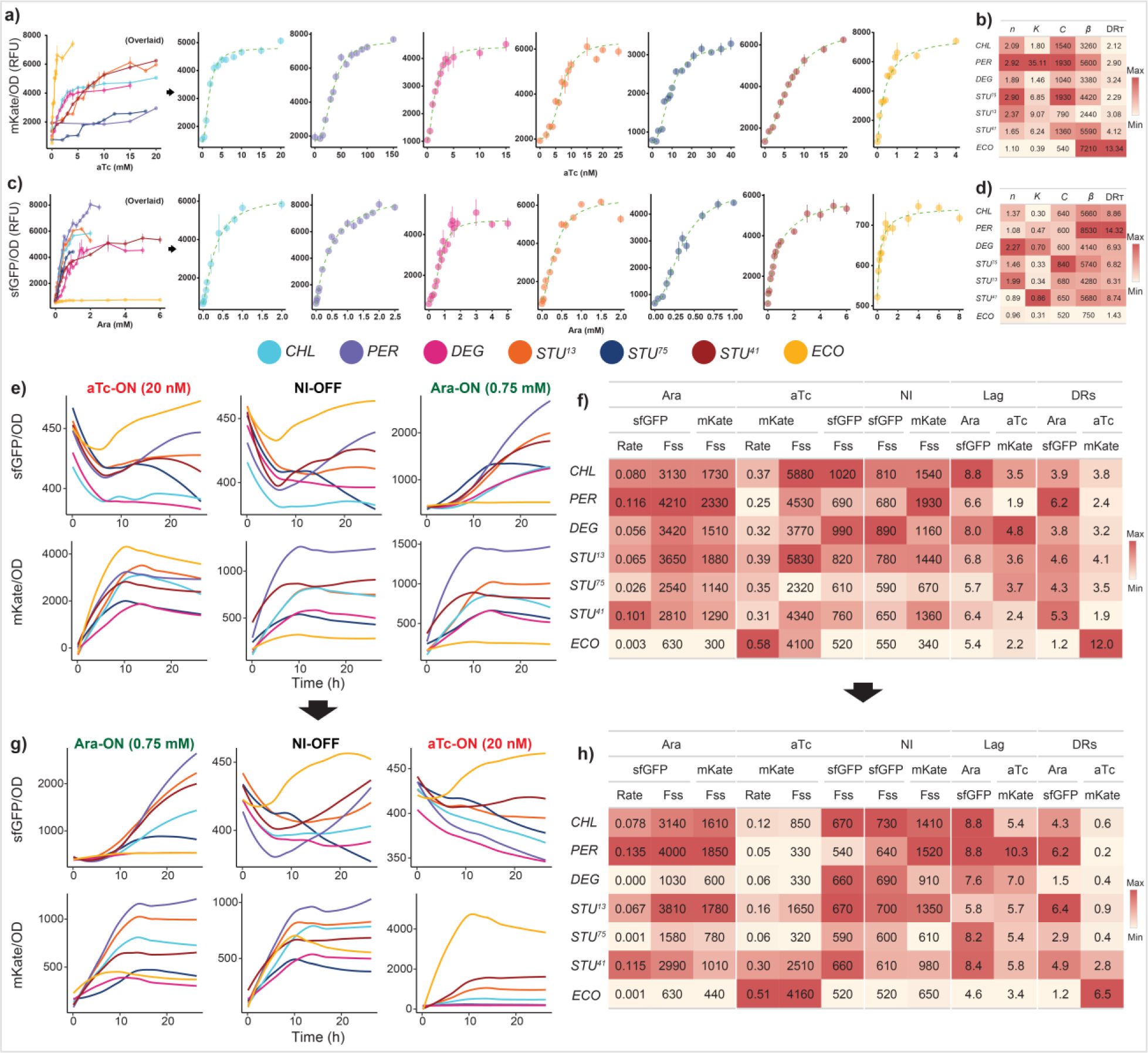
The chassis-effect is observed through the measurable performance of the genetic inverter between closely related *Stutzerimonas* hosts. a) aTc induction curves with left-most plot showing overlaid induction curves up to given inducer concentration, followed by individual induction curves with scaled axis. Hosts are color coded, error bars indicate standard error of the mean, n = 8. b) Estimated Hill parameters from aTc induction curves. Color scale relative to each column. c) Ara induction curves and d) estimated Hill parameters. e) OD_600_ normalized fluorescence dynamics of one of three toggle assays with induction scheme 0.75 mM Ara and 20 nM aTc. Initial OFF cells were diluted to respective induction states. f) Estimated fluorescence metrics from fluorescence in e) across induction state and fluorescence output type. g) Fluorescence dynamics of toggled cells diluted to respective opposite inducer and h) estimated fluorescence metrics. NI = no induction, Fss = late phase steady-state fluorescence, Rate = max specific rate. DRs = Specific dynamic range.

We next performed a toggle assay to demonstrate the invertibility of the device and to further quantify the chassis-effect by the different degrees of hysteresis experienced by each host (Fig. 2e-h). Initial OFF cells (grown in absence of inducer) were diluted into media with inducer to prompt Ara-ON or aTc-ON states, respectively (Fig. 2e) and performance metrics from normalized sfGFP and mKate fluorescence curves was determined across induction states. These metrics (Fig. 2f) include the maximum steady/state fluorescence at late growth phase (F_ss_), maximum rate of fluorescence (Rate) at exponential phase, specific dynamic range (DR_S_) and lag time (Lag). Each of these metrics have different biological implications, depending on the induction state. For example, the fluorescence output under OFF conditions is a measure of the inverter’s unbiased output level – i.e., the host-specific background fluorescence. Meanwhile, fluorescence outputs from cells grown under presence of an antagonistic inducer (e.g., sfGFP output in presence of aTc) indicate the deficient transcriptional control (leakage). The range of observed fluorescence curves and estimated performance metrics further solidifies the existence of the chassis-effect. Inversion between states was controlled by washing cells twice and diluting them into to media containing respective opposite inducer (Fig. 2g, h).

Results from the toggle assay showed that upon being toggled from Ara-ON to aTc-ON, multiple hosts experienced an attenuated mKate output, to different degrees. For instance, the DR_S_aTc_ values of all hosts other than *S. stutzeri* DSM 4166 all drop below 1 when diluted from Ara-ON to aTc-ON, meaning their induced mKate output becomes lower than their baseline output. This decreased output was not observed when diluted from OFF to aTc-ON and can therefore be attributed as a measurable hysteresis effect. Two additional toggle assays with induction schemes 0.25 mM – 40 nM aTc and 0.375 mM Ara – 5 nM aTc was performed (Figure S1), and for *S. chloritidismutans*, *S. degradans* and *S. stutzeri* DSM4166, the mKate output attenuation was relieved under induction schemes with decreased Ara concentration but hosts the other three hosts experienced consistent attenuation across toggling schemes. Since cells were washed twice and diluted, the mechanism in which mKate output is attenuated is likely not due to residual extracellular Ara concentrations. This result suggests the invertibility (input/output logic) of the inverter is dependent on past induction states and that dynamics of this hysteresis-effect varies between hosts based on intracellular molecular physiology.

### Species-specific physiology responses to the genetic inverter

A consistent growth inhibition was observed when comparing wild type hosts against their engineered genotypic counterparts and across induction states (Fig. 3). Growth of hosts operating the genetic inverter was measured simultaneously during the toggle assay from which growth physiology was characterized (Fig. 3a-d). The reference host (*E. coli*) demonstrated an overall more rapid growth on LB media compared to all *Stutzerimonas* hosts. The addition of BB23 backbone lead to decreased specific growth rates of all strains compared to their respective wild type counterparts, with *S. stutzeri* 24a75 exhibiting the most reduced growth rate along with *S. stutzeri* 24a13 and *E. coli*. (Fig. 3e, f). This is expected due to the added burden of maintaining the vector backbone in presence of kanamycin. *E. coli* and *S. chloritidismutans* exhibits the greatest additional growth burden upon addition of the inverter device into the backbone (complete pS5 plasmid) under NI conditions. The added burden is likely due to larger plasmid payload^32^ and possible leakage expression of the device. However, some hosts experience close to zero additional growth burden, and while *S. chloritidismutans* had relatively high amount of leakage expression (sfGFP and mKate output in absence of inducer) (Fig. 2c, right), *E. coli* exhibited the lowest leakage expression, suggesting the mechanism in which the backbone vector and device imposes growth inhibition on its host varies between hosts. Induction of the genetic inverter exacerbated growth inhibition (Fig. 3e, f), which also varied within hosts. For instance, the growth rate of *S. stutzeri* 24a75 was almost halved when induced with Ara when compared with NI condition but showed little response when treated with aTc. Overall, these results exemplifies that the genetic inverteŕs mode of operation uniquely affects the growth of each host.

**Figure 3.**
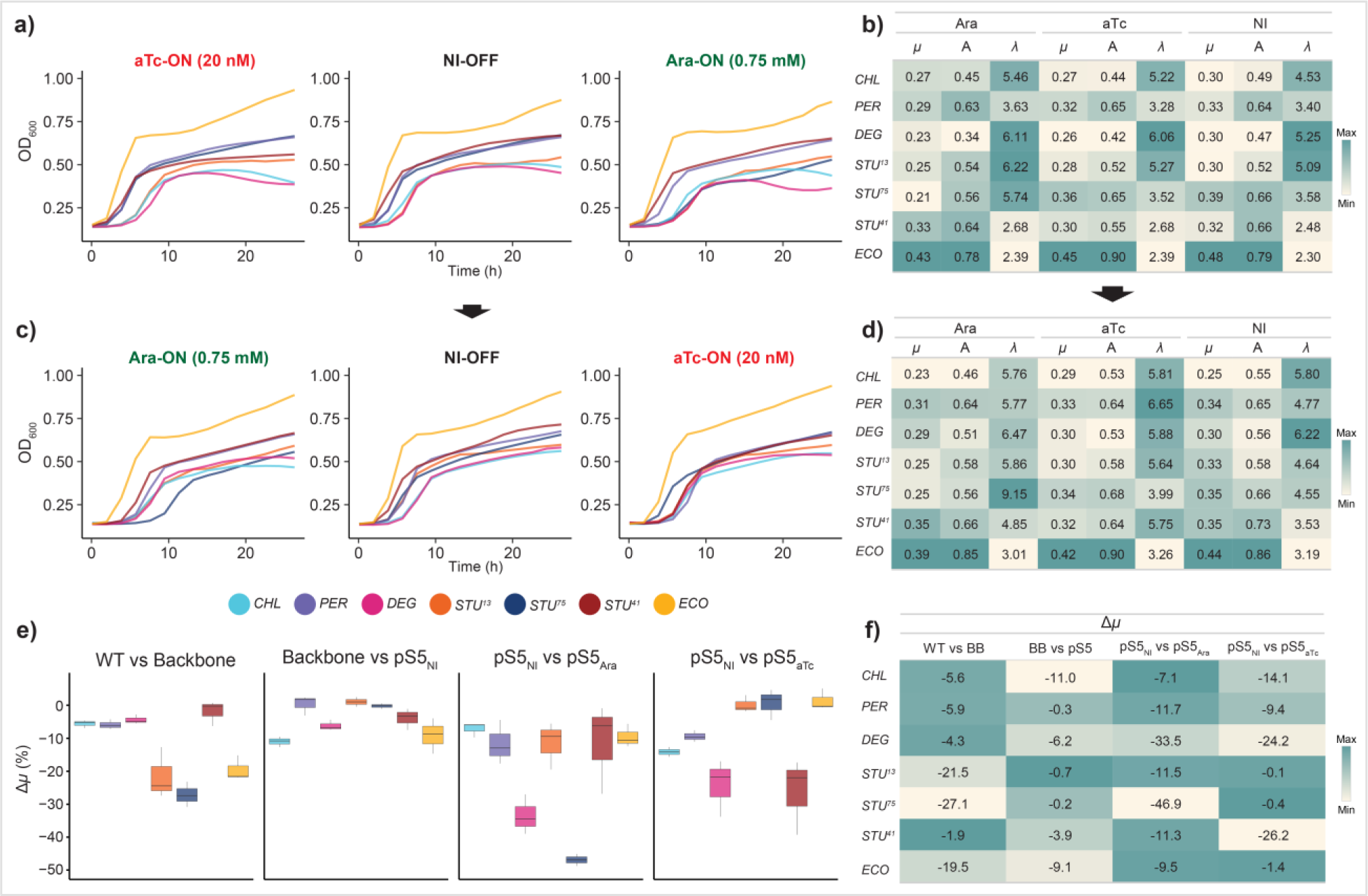
The growth dynamics are uniquely affected as a result of host-specific operation of genetic inverter. a) Growth curves of initial OFF cells diluted to respective induction states. Hosts are color coded. b) Estimated growth metrics for each host from growth curves in a). Color scale is relative to each column. c) Growth curves of toggled cells diluted to respective opposite inducer to toggle induction state and d) corresponding estimated growth metrics. e) Growth difference between host genotypes and/or induction state captures by the Δ*µ* metric, defined as relative percentage change in growth rate. WT = wild type, BB23 = pBBR1 and KanR cloning vector. pS5_NI_ = pS5 plasmid in absence of inducer. pS5_Ara_ = pS5 plasmid in presence of Ara (0.75 mM), pS5_aTc_ = pS5 plasmid in presence of aTc (20 nM). NI = no induction; *µ* = max specific growth rate; A = carrying capacity; *λ* = lag time.

The unique intracellular molecular environment within each host is ultimately shaped by their respective genomes, and perhaps more importantly, the expression pattern of their gene products. We therefore extracted total RNA from cells induced in both directions for mRNA sequencing to determine their transcriptome profiles. By augmenting our comparative transcriptome analysis with pangenomic insight, we can discern the impact that the core and accessory genome has on the observed host-specific inverter performance.

### Unique transcriptional patterns arise from both presence and operation of an engineered genetic circuit

Global transcriptome analysis revealed marked variability in gene expression profiles between hosts operating the engineered genetic inverter. The difference in response was observed in terms of magnitude (number of differentially expressed genes or DEGs), the functional composition of DEGs and direction of regulation of certain gene clusters shared among hostś core genomes. These unique transcriptomic profiles support our hypothesis that differences in gene expression from core and/or accessory genomes uniquely correspond with differences in genetic inverter performances.

Cross-species comparison of DEG profiles was performed by pooling all read counts mapped to gene calls within a gene cluster, on the basis that genes within the same gene cluster are inferred to be highly similar. Differential expression of mRNA encoded from the genetic inverter confirmed its programed operability (Fig. 4). Cells of each host that were treated with Ara showed a higher proportion of reads (measured by transcripts per million, TPM) mapped to *tetR* and *sfGFP* compared to *araC* and *mKate,* and vice versa for aTc induced cells (Figure S2). Differential gene expression analysis of Ara against aTc treated hosts reveals significant upregulation of *tetR* and *sfGFP* genes and downregulation of *araC* and *mKate*, in accordance with the design of the inverter. The degree of response differed greatly between hosts. Log2 fold-change values of *tetR* ranged from 4.9 (*S. chloritidismutans*) to 8.2 (*S. degradans*) and ranged from 1.1 (*S. perfectomarina*) to 4.1 (*S. stutzeri* 24a75) for *sfGFP*. These results support the observed chassis-effect via mRNA abundances from genes encoded within the engineered genetic inverter. The consistent downregulation of the *kanR* gene in Ara induced cells could be due to higher expression of *kanR* in aTc induced cells because of transcriptional readthrough when transcribing from the pTet promoter. A relatively large difference in TPM values was also observed between polycistronic repressor-reporter pairs (Figure S2).

**Figure 4.**
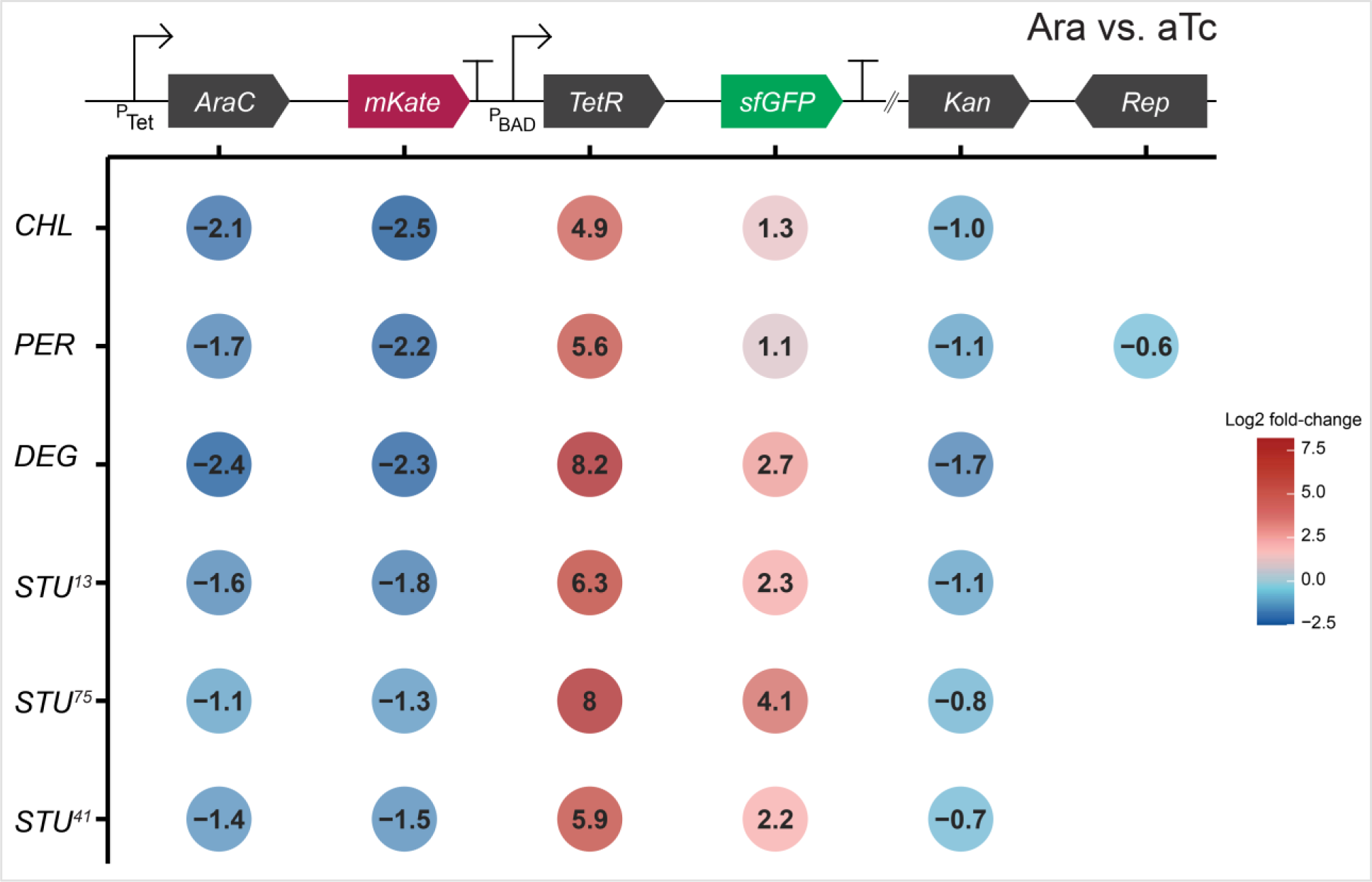
The chassis-effect is measurable at the differential response of genes designed into the engineered genetic inverter. Log2 fold-change values of the six genes encoded in the pS5 plasmid between hosts comparing Ara against aTc induced cells. Empty position indicates non-significant differential expression.

The mRNA abundance of genes encoded by the genetic inverter intuitively play a major role in influencing the observed chassis-effect, but these heterologous gene products are not compartmentalized away from native cellular elements. We were therefore prompted to also explore the hosts’ global transcriptome response to better understand how expression of genes from the core and accessory genomes might be concordant with the chassis-effect. Our results reveal a clear diversity in the global differential gene expression responses between hosts when comparing Ara against aTc induced cells, indicating that genome encoded functions respond differently depending on how the engineered genetic inverter is being operated (Fig. 5a, b). Differential expression analysis with a *P*-value threshold of 0.05 and log2 fold-change threshold of 0 leads to a total of 4,825 significantly differentially expressed genes (DEGs) distributed across hosts, with 3,995 DEGs belonging to the core group distributed among 1672 gene clusters. We observed that each host significantly expressed only a subset of shared core gene clusters, with 63 % of core gene clusters being expressed by at least three hosts. Among significant DEGs, the direction of regulation and the strength of response (captured by log2 fold-change value) within a gene cluster differs (Fig. 5c). The number of DEGs is unevenly distributed between the hosts, varying from 589 (*S. chloritidismutans*) to 1106 (*S. stutzeri* 24a75) (Fig. 5d) meaning the operation of the genetic inverter (*i.e.,* the user defined induction state) incites a greater response in some hosts than others. We note that the differences in DEG response between states cannot be entirely attributed to the expression of the inverter alone, but also from the presence of inducer compounds Ara and aTc. However, the addition of induction compounds is inherently coupled to the function of the inverter by design and can therefore include responses to inducers as a contributor to the chassis-effect.

**Figure 5.**
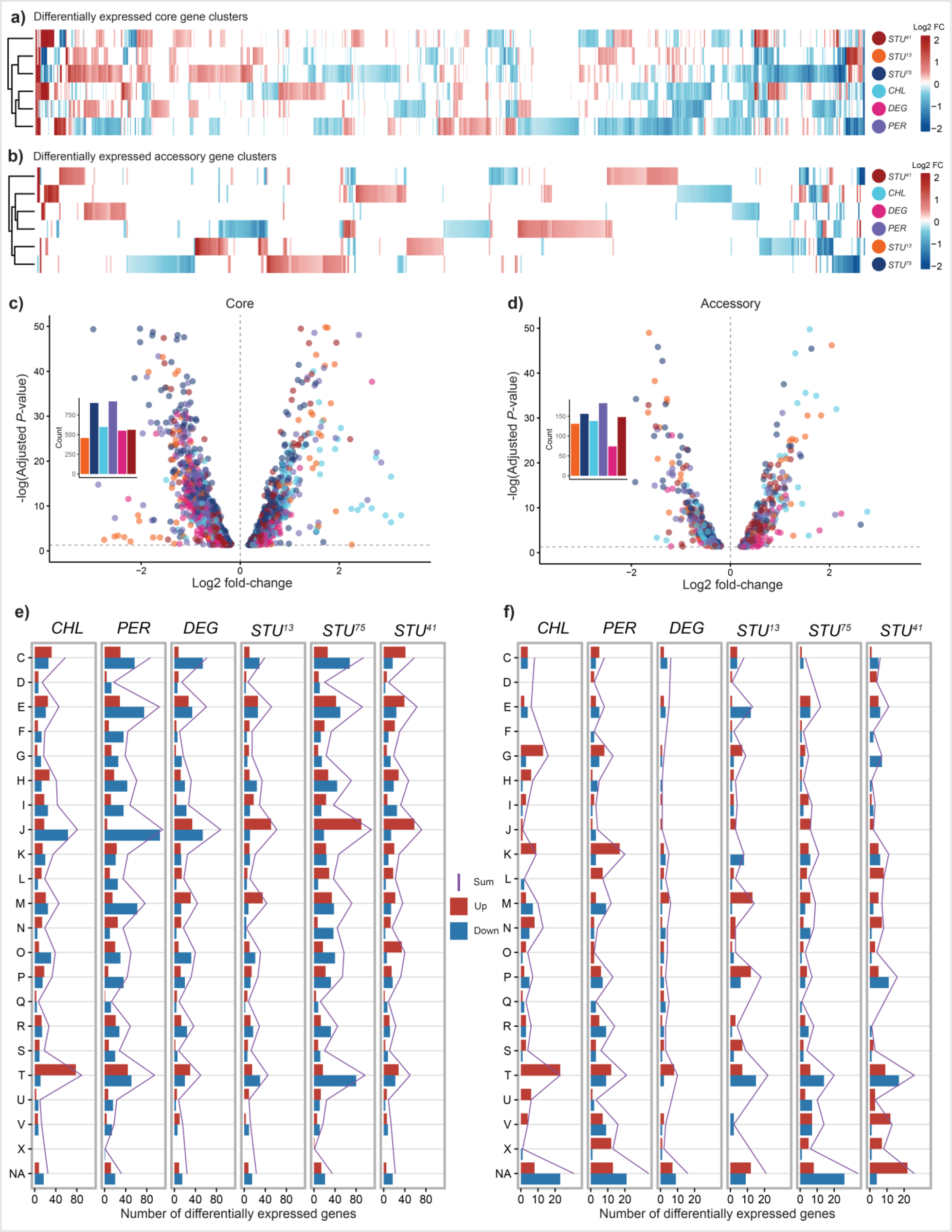
Global differential gene expression analysis of Ara against aTc induced cells reveals diverse transcriptional profiles as a result of inverter operation. Log2 fold-change values of differential expressed a) core and b) accessory gene clusters significantly expressed by at least one host (*P*-value < 0.05). White bar indicates non-significant expression in host. For accessory genome, white bar indicates either non-significance or gene cluster has no hits for that host. Volcano plots visualizing log2 fold-change distribution of significantly DEGs in c) core genome and d) accessory genome. Inset bar charts show the number of differentially expressed genes for each host. The distribution of DEGs across COG categories across hosts are shown for both e) core and f) accessory. DEGs are further grouped into upregulated (red bars) and downregulated (blue bar) gene clusters within each COG category. The purple line denotes the sum of number of DEGs.

We next examined the functional profile of the core and accessory DEGs by assessing their distribution among the COG categories. The core DEGs of hosts shows a surprisingly uniform distribution across the 22 COG categories while a more varied response pattern in the accessory genome is observed. 45.3 ± 1.5 % of the DEGs can be found in categories C, E, J, M, T alone (Fig. 5e). Meanwhile in the accessory genome, COG categories P, K, T and NA (unassigned) make up 43.0 ± 2.6 % of DEGs (Fig. 5f), with NA gene clusters making up 17.1 ± 2.3 % alone. We observed difference in the composition of up and down regulated genes between hosts within functional groups (especially J, T, and M in the core group) suggesting varied responses to inverter activity. The less uniform transcriptomic responses from accessory genomes are attributed to gene clusters shared by only a subset of species. In the accessory genome, larger numbers of DEGs were concentrated in category T (Signal Transduction) for all hosts, indicating unique response patterns occurring within each host as a result of inverter activity. A larger variation in distribution of DEGs was observed within a COG category as well. For instance, 31.3 % of DEGs in category K occured in *S. perfectomarina* alone, further exemplifying the diversity in gene expression response pattern. We. note that the majority of DEGs are unassigned (NA), meaning with improved gene annotation, the transcriptional profile of the accessory DEGs could change substantially. The accessory genome, comprising genes with specialized functions shared by only a subset, can be a strong source for genes contributing to an observed chassis-effect. However, differential expression of functions encoded within the core genome often belong to the major carbon, nitrogen and energy metabolism, cell division and housekeeping genes, which can more intuitively underpin the chassis-effect, given that majority of DEGs stem from the core genome.

### Differential expression of core genes is concordant with the chassis-effect

Procrustes Superimposition (PS) analysis revealed significant concordance between the host specific expression of the core genome and genetic inverter performances. In contrast, there were no significant correlations between the accessory genome and device performances. Hence, the genome encoded functions most responsible for the observed chassis-effect arise from genes shared commonly between hosts. PS analysis compares the similarity between two configurations on a coordinate, and when the distances between points the configuration carry information of (dis)similarity, such as points on principal components, PS analysis can be used to determine whether similarities between two datasets correlate^33,34^. PS analysis was therefore performed to formally test for concordance between the PCA ordinated configurations of the hosts in terms of their similarity of genetic inverter performance and expression of their genome. This data exploration was done in a hierarchal manner, by first considering all significant DEGs, and then dissecting the transcriptome into core and accessory. Significancant correlations were measured when comparing the configurations from inverter performance and when taking into consideration all DEGs (Fig. 6a, *P*-value = 0.044, m^2^ = 0.344). Upon splitting the DEGs into respective core and accessory groups, we found that it is in fact the core genome that is responsible for the significance observed previously (*P*-value = 0.034, m^2^ = 0.324) (Fig. 6b, c). This result supports the following notions, that the observed chassis-effect can be explained by the differential gene expression response between hosts and that hosts with more similar expression pattern of their shared core genome also have significantly more similar performances. Hence, our given set of species and their shared gene clusters, core genes commonly associated with housekeeping functions, central carbon, and energy metabolism, are major biological drivers of the chassis-effect. Additionally, PS analysis applied to the captured growth metrics show significant correlation with inverter device performance (Figure S3), corroborating our previous result that observed differences in the physiological state of hosts can be used to potentially predict device performance^14^.

**Figure 6.**
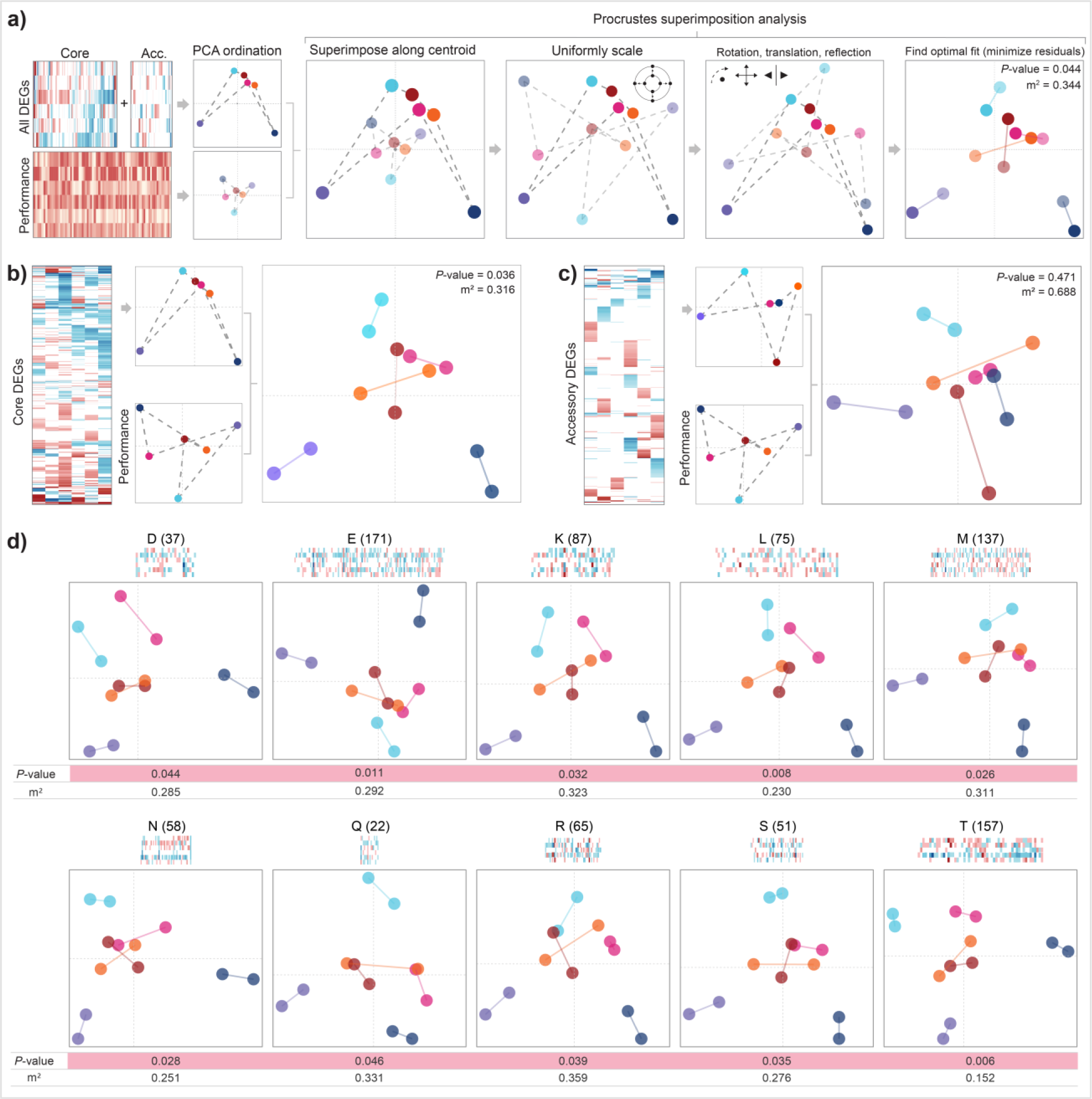
Procrustes analysis reveals significant concordance between similarity in inverter performance and similarity in core genome response between hosts. a) Procrustes superimposition analysis comparing hosts in terms of their total significant differential gene expression response against inverter performance metrics. The key steps in PS analysis are schematically illustrated. Performance metric dataset and differential gene expression dataset are first projected onto ordinate space via PCA, then the configurations are compared through PS analysis, which involves centering, scaling, and transforming the two projections with the goal of minimizing the sum of squared vector residuals (the m^2^ statistic) between each respective point (host). Significance of the obtained statistic is determined through a permutation method. Dashed gray lines between points have been added to visualize an arbitrary “configuration” formed by each dataset, which connects each point in the following arbitrary order “-*CHL*-PER-*DEF*-*STU*^13^-*STU*^75^-*STU*^41^-”. b) Procrustes analysis comparing inverter performance against significantly differentially expressed b) core and c) accessory genome. d) Sub-groups of core genome grouped by COG category that were significantly correlated against inverter performance as determined by PS. Number in sub-plot headers indicates number of genes clusters within each group along with log2 fold-change profile of gene clusters. For gene clusters assigned to multiple COG categories, it was counted as belonging to each of those COG categories.

We next interrogated the specific cellular functions that are most concordant with the observed chassis-effect by dissecting the differentially expressed core and accessory genomes into their functional categories in a second iteration of Procrustes analysis. Ten of the twenty-three COG groups were found to be significant (Fig. 6d). Category E (Amino acid Transport and Metabolism) is especially intuitive because the main carbon and energy source provided from LB media are amino acids (not sugars) and different catabolic strategies for these will in turn cause different growth phenotypes. Categories R (General Function Prediction Only) and S (Unknown Function) were also significant, suggesting that genetic elements of unknown function are contributing strongly to the observed chassis-effect.

### Genes clusters involved in denitrification and efflux pumps are highly responsive to genetic inverter activity within *Stutzerimonas* spp

A data-driven investigation of the top 100 most differentiated gene clusters between host cells revealed that gene clusters involved in denitrification, iron acquisition and membrane-bound transport proteins are among the most highly differentially expressed gene clusters in response between Ara versus aTc treatment applied to the *Stutzerimonas* hosts (Fig. 7). This was determined by ranking gene clusters by a metric that takes into consideration the number of hosts significantly expressing each gene cluster, the sum of absolute log2 fold-change values and the combinatorial sum of absolute differences in log2 fold-change values between species (Table S1). To add confidence in the functional annotation of the gene clusters, we supplemented the COG annotation with annotation using the KEGG Orthology (KO) database^35^ because COG accessions are (in some cases) only general descriptions of protein functions or families. For instance, the assigned COG accession for the four highly ranked gene clusters IGC_00001064, GC_00001456, GC_00001627 and GC_00001727 describes all these gene clusters to encode for cytochrome c protein (CccA, COG 2010). The corresponding KO annotations for the four *CccA*-annotated gene clusters are cytochrome c55X (NirC, K19344), dihydro-heme d1 dehydrogenase (NirN, K24867), nitrite reductase (NirS, K15864) and cytochrome-containing nitric oxide reductase subunit c (NorC, K02305), respectively, which are all heme-containing enzymes belonging in the cytochrome c protein family^36,37^. Notably, every KO accession has a complementary COG accession entry in the KEGG Orthology database (https://www.genome.jp/kegg/ko.html), and the KO and COG annotations done independently by KofamKOALA and Anvi’o (for COG annotation) discussed here all match their respective database records, indicating that the COG and KO annotations are in agreement.

**Figure 7.**
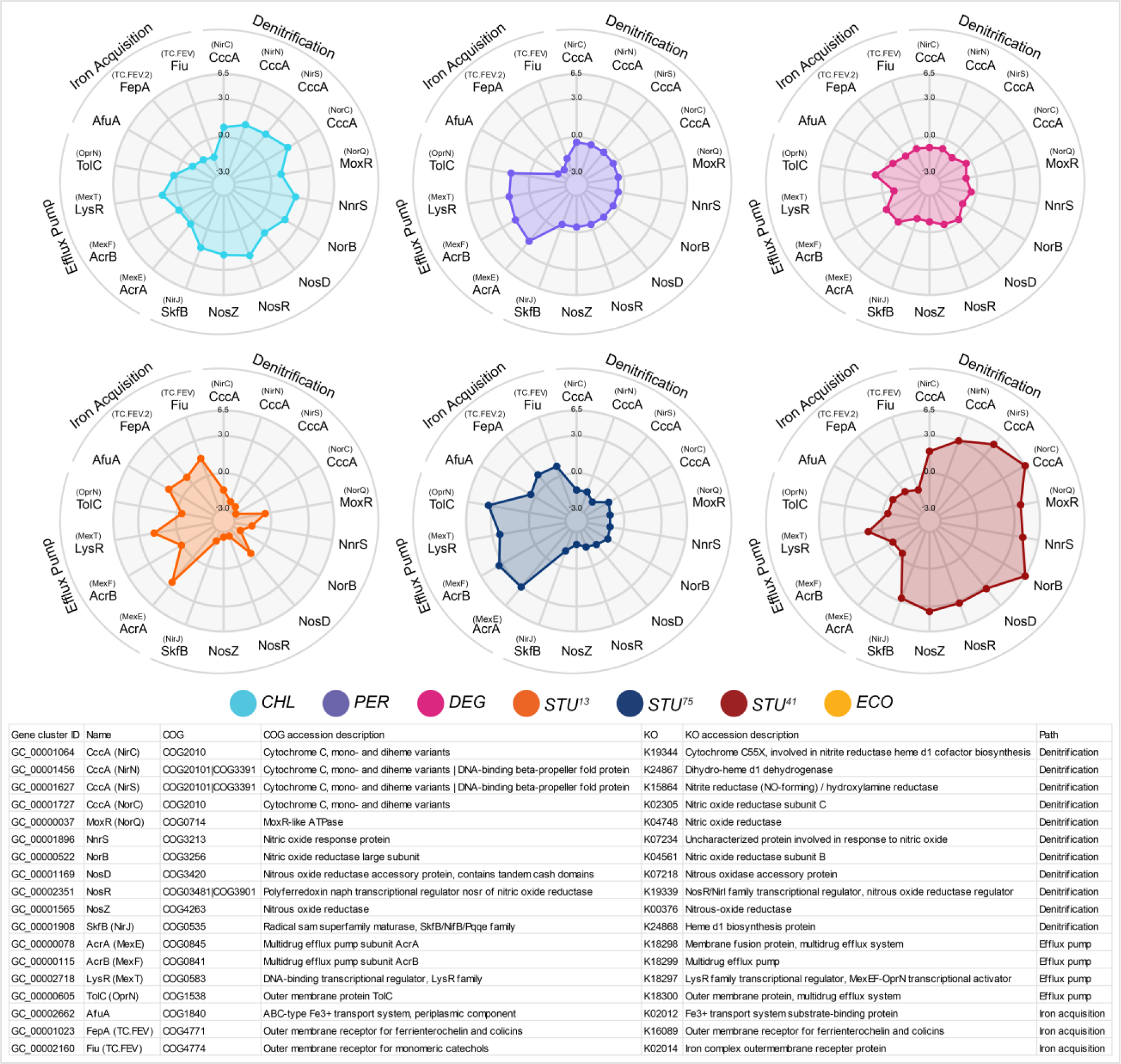
Gene clusters containing the most highly differentially expressed genes between hosts. Spider plots showing log2 fold-change data of most highly differentially expressed gene clusters between the hosts. Gene names in parentheses are names provided by KO annotation, all other gene names are provided by the annotated COG accession. In cases where the gene name provided by COG and KO match, only one gene name is shown.

Genes involved in denitrification were upregulated in *S. chloritidismutans* and *S. stutzeri* DSM 4166 and downregulated in *S. stutzeri* 24a13 and *S. stutzeri* 24a75 but showed little change in *S. decontaminans* and *S. degradans* when comparing Ara against aTc induction states. This result was not anticipated and exemplifies that operation of engineered genetic devices can impact and/or be impacted by fundamental cellular functions seemingly unrelated to the heterologous transcription and translation networks. Denitrification is canonically described as an anaerobic process, but aerobic denitrification has been identified in numerous bacteria, many of whom have been identified as *Pseudomonas stutzeri* species^21,38,39^. The gene *napA* encodes for a nitrate reductase which catalyzes the important first step in the denitrification pathway. Only one gene cluster, GC_00001817, was annotated as *napA* by KO annotation, but this gene cluster was only significantly differentially expressed in *S. decontaminans* and ranked low. Genes involved in iron acquisition showed the opposite trend, being upregulated in *S. stutzeri* 24a13 and *S. stutzeri* 24a75 and downregulated to some degree in all other hosts. Denitrification activity requires bioavailable iron for heme biosynthesis^40^, hence we expected the differential expression of the two pathways to instead positively correlate.

Other highly ranked gene clusters of note from the data-driven investigation are GC_00002718, GC_00000078 and GC_00000115. COG annotation inferred these gene clusters to encode for components of the AcrAB-TolC efflux pump. Meanwhile, KO infers them to be components of the MexEF-OprN efflux pump. Both pumps are broad-substrate Resistance-Nodulation-Division family transporters known to provide drug resistance^41,42^. Another high-ranking gene cluster was GC_00002718, annotated by COG as a transcriptional regulator part of the LysR family, but as MexT by KO, the latter being a LysR-type transcriptional regulator regulating the MexEF-OprN operon. The inclusion of the LysR regulator indicates that gene clusters GC_00002718, GC_00000078 and GC_00000115 indeed encode for a MexEF-OprN efflux pump, as AcrAB-TolC is regulated by the transcription factor AcrR, which instead belongs in the TetR family of transcriptional regulators^43^. Interestingly, hosts that exhibit downregulated or low change in expression of denitrification gene clusters were also the hosts with highest upregulation of the AcrAB-TolC/MexEF-OprN encoding gene clusters (hosts *S. decontaminans*, *S. stutzeri* 24a13 and *S. stutzeri* 24a75). Incidentally, Fetar, et al. 2011 has reported that nitrosative stress in the form nitric oxide accumulation is a direct inducer of MexEF-oprN expression^42^, suggesting that hosts expressing the efflux pump could be a response to nitric oxide.

## DISCUSSION

This study was driven by the overarching question as to whether closely related bacterial hosts can exhibit a strong chassis-effect when programmed with an identical engineered genetic inverter and how contributions from the core and accessory genomes might underpin this phenomenon. This was investigated using a supervised machine learning approach that employed dimension-reduction techniques combined with multivariate statistical analysis to determine which portions of the differentially expressed genome were concordant with the measured chassis-effect and then further trace these differences into specific genome encoded functions. In addition to our main finding, we report the successful engineering of several *Stutzerimonas* species. *S. degradans* is of interest with its known applications in bioremediation of contaminants^18^ and *S. stutzeri* 24s75 is inferred to possess the *nifDKH* genes according to KEGG pathway mapping^44^ and therefore has potential application as a biological nitrogen fixation agent to replace agrochemicals, similar to *S. stutzeri* DSM 4166, the latter in which has previously received much attention for its nitrogen fixation capability^27^.

It is tempting to theorize that accessory and host-specific unique genes would be the leading cause of phenotypic distinction (and resulting chassis-effects), yet our findings empirically showed no such relations between inverter performance and the accessory genome. Instead, we observed concordance between transcriptional response of shared core genes and inverter device performance. In other words, hosts with more similar gene expression from their core genomes also have more similar performance, suggesting that the expression of the core genome, and in turn its protein products, have a greater impact on the cellular state. This holds especially true if the accessory genome consists of genes that are not expressed unless conditions call upon their expression, such as for unique biosynthetic gene clusters^45^. The core genome of our *Stutzerimonas* platform is enriched for genes involved in central carbon and energy metabolism as well as housekeeping-associated functions. Combined with the result that observed differential growth physiology between hosts was found significantly correlated with differential performance, the chassis-effect may manifest through the redistributive flux of cellular resources and gene expression machinery unique to each host context because of genetic device maintenance and operation. This concurs with previous studies that has demonstrated how resource dynamics such as ribosome abundance^46^ can affect genetic circuit performance and even parameterized resource competition to build predictive models^47,48^.

Data-driven analyses showed that denitrification genes were responsive to heterologous expression of the engineered inverter. This indicates that engineered gene circuits can have unforeseeable and unspecific cross talk with fundamental cellular processes and that these events are context specific to individual hosts. If denitrification is in fact occurring, the source of nitrate in the media is unclear and would require a more targeted analysis on proteins and metabolic intermediates involved in denitrification pathways. A possible explanation is that our hosts can perform nitrification of ammonia to produce nitrate, with the source of ammonia being the by-product of amino acid catabolization from the oligopeptides in the LB media. But KEGG pathway mapping reveals that neither of two the genes involved in the canonical nitrification pathway (ammonia monooxygenase, *amo*, and hydroxylamine reductase, *hao*) are found in the genomes of our hosts (Supplementary Figure S4), suggesting a lack of nitrification capability. However, there has been numerous reports of other *Stutzerimonas* (reported as *Pseudomonas stutzeri*, now deprecated) capable of converting ammonia to nitrogen gas through heterotrophic nitrification and aerobic denitrification (HNAD) despite their genomes encoding neither *amo* nor *hao*^49–52^. These reported HNAD capable species are thought to perform HNAD through an undocumented pathway, or the enzymes catalyzing the reactions performed by *amo* and *hao* are significantly different enough in amino acid composition to escape current annotation algorithms. The same enigmatic HNAD process could be occurring for our *Stutzerimonas* species with increased denitrification activity. Denitrification a pragmatic phenotype due to its bioremediation potential^53^, but bacteria capable of simultaneous HNAD are valuable as the cost of maintaining anaerobic conditions is reduced^21^.

Given the complex interwoven network structure of cellular metabolism^54^, it is impractical to postulate that the observable chassis-effect between a given set of hosts can be explained by a single or even a set of predictable genome-encoded functions without experimental insight. The examples from this study were dentification and efflux pumps, but given that it is host-context-specific, device operations in other species/strains may influence different cellular functions. Hence, deriving a mechanistic metabolic model detailing how these factors interact to bring about the produced output is likely intractable and may not hold true across different sets of hosts. Incidentally, with the advent of dimension reduction using multivariate statistical models in supervised machine learning, synthetic biologists can advance data-driven engineering strategies and bypass dependence on *a priori* mechanistic insight for biodesign applications^55,56^. In contemporary biological research, unveiling a mechanistic model that explains an observation is considered the pinnacle goal, but the goals of synthetic biologists more often prioritize the practical implementation of designed systems rather than knowledge-acquisition. Supervised machine learning aligns well with this goal^57^.

We have developed new knowledge of how observable chassis-effects of an engineered microbial platform can be explained by genome encoded functions that are largely conserved and represent fundamental cellular processes. In addition, we uncovered a context-specific phenomenon that engineered genetic devices have unpredictable interference with metabolic functions that regulate cellular physiology. The evidence for these conclusions is corroborated by changes in species and even strain specific global transcriptome profiles and growth phenotypes, and the use of multivariate statistics to attribute how quantifiable chassis-effects are concordant with genome structure and function. This represents a genome-informed advancement within the field of broad-host-range biodesign which aims to lessen our reliance on model organisms so that we can better understand diverse microbial behaviors and use them in the blueprints of biodesign. The number of assimilated microbes available for use as industrial biotechnology platforms and biodesign engineering is increasing steadily^58–60^, heralding the advancement towards broad-host-range synthetic biology where synthetic biologists must explore not only the design space of genetic parts, but also the design space of host-chassis in order to optimize their engineered systems. As the field of biodesign progresses towards this new era, the constraints imposed by the chassis-effect will only become more relevant. It is therefore of high interest to develop novel hosts and data-driven predictive frameworks capable of using genomic insight to understand how chassis-effects might limit or in some cases expedite design-build-test cycles for the futurés biotechnology applications such as bioremediation of soil and marine contaminates and production of renewable agrochemicals.

## MATERIALS AND METHODS

### Species, cultivation, cloning, and transformation

Overview of species used in this study can be found in Table S2. The six *Stutzerimonas* hosts selected for further study were identified by verifying *rpoD* sequence through sanger sequencing^61^ (Table S3). Cells were cultured in Lysogeny-Broth (LB) at 35 °C unless specified otherwise. BB23 backbone and pS5-carrying strains were cultivated in presence of 100 µg/mL kanamycin while wild types were grown without. Single colonies from streaked LB agar plates were picked to inoculate liquid media and incubated overnight with shaking to prepare overnight cultures. 199 µL of media was inoculated with 1 µL of overnight culture in black clear-bottom 96-well plates (Thermo Fischer, 165305) and sealed with Breath-Easy film (Sigma-Aldrich, Z380059). OD_600_, sfGFP (Ex 485/ Em 515, gain 75) and mKate (Ex 585/ Em 615, gain 125) fluorescence was measured continuously using a Synergy H1 plate reader (Agilent Biotek, Serial Number 21031715) with continuous linear shaking (1096 cpm, 1 mm) at 9 mm read height. Working stock solutions of 1 M L-Arabinose (VWR, A11921) stock and 1 mM aTc (VWR, CAYM10009542) was prepared by dissolving powder in water and 70% ethanol respectively. Cloning was performed using *E. coli* DH5α, made chemically competent and transformed following the Inoue method^62^. *Stutzerimonas* species were transformed via electroporation method as previously described in Chan et al. 2023^14^. Primers used in this study can be found in Table S4. BB23 backbone was integrated into the BASIC Assembly format using from pSEVA231 as template with primers B_SEVA_F and B_SEVA_R. Plasmid pS5 was assembled in the Biopart Assembly Standard for Idempotent Cloning (BASIC)^63,64^ environment as previously described^14^. Sequence and accession of pS5 components can be found in Table S5.

### Induction assays

Overnight culture grown in absence of inducer was used to inoculated to media with various concentrations of aTc and Ara in 96-well plates. The normalized steady-state fluorescence at late growth phase (Fss) averaged oved a time window of 6 – 12 hours was used as response variable of induction curves. In R (v4.3.1), Hill coefficient (*n*), activation coefficient (*K*) and max steady-state fluorescence output (*β*) was estimated by fitting the Hill function (1) using non-linear least-square regression with the “nls” function from the stats base R package. For parameter *C*, representing basal fluorescence output at 0 inducer concentration, empirical value was used.

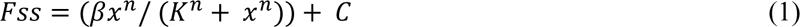

Where x is Ara (mM) or aTc (nM) inducer concentration.

### Toggle and growth assay

Overnight culture grown in absence of inducer was used to inoculate media in 96-well plates supplemented with Ara, aTc and no inducer condition. To toggle, cells were harvested by centrifugation at 4000 RPM for 20 minutes at room temperature and supernatant removed before resuspending in 200 µL LB media, this washing step was repeated for a total of two washes. After final resuspension, 1 µL of washed cells were inoculated to 199 µL fresh media supplemented with the opposite respective inducer. Growth difference metric Δ*µ* was calculated using equation 2.

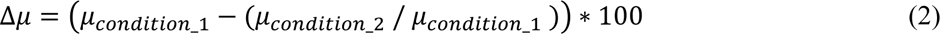

Where *µ* is max specific growth rate and “condition_1” and “condition_2” denotes sample condition in terms of genotype and induction state. In R, max rates of OD_600_ and normalized fluorescence curves were estimated based on a rolling regression method using the “all_easylinear” function from the growthrates (v.0.8.4, https://CRAN.R-project.org/package=growthrates) R package. Lag times and curve plateaus of OD_600_ and normalized fluorescence curves were determined using the “all_growthmodels” function, fitting the Gompertz growth model^65^ with additional lag (*λ*) parameter.

### Pangenome analysis

Genomes of *Stutzerimonas* hosts was downloaded from NCBI. Comparative pangenomic analysis was performed in the Anvi’o (v7.1) environment. pS5 plasmid nucleotide sequence was manually added to each *Stutzerimonas* genome file. For complete data on binned gene clusters obtained comparative pangenomic analysis, see Data and Code Availability. Unless specified, all Anvi’o commands were run using default settings. Briefly, genomes were converted to contigs databases using “anvi-gen-genomes-storage” command, which uses Prodigal^66^ to make gene calls. Gene calls were annotated with the Clusters of Orthologous Genes 2020^67^ database and KEGG Orthology^35^ database. Pangenome analysis was done using “anvi-pan-genome” command with --mcl-inflation parameter set to 10 for high cluster granularity and DIAMOND was run with “-sensitive” flag as recommended when comparing genomes from closely related organisms. Gene clusters were binned according to their number of occurrences across the six genomes in the interactive Anvi’o pangenome display command “anvi-display-pan”, from “Bin 1” to “Bin 6”. We define gene clusters as belonging to the core genome if it has hits in all six genomes (Bin 6). All other gene clusters were defined as accessory, with gene clusters with only one hit (“Bin 1”) further distinguished as unique.

### Total RNA extraction and RNA sequencing

Cultures for total RNA harvesting was initiated as described for toggle assay with induction scheme 0.75 mM Ara and 20 nM aTc. Cells were harvested at late exponential growth phase by centrifugation at 10 000 rpm for 1 minute before discarding the supernatant and immediately freezing in liquid nitrogen. Total RNA was extracted using the Quick-RNA Miniprep Kit (Zymo Research, R1055) following manufacturer’s instructions. The kit includes a cell lysis step and 15 min on-column DNase I treatment at room temperature. RNA samples in triplicates for each group were sent to Eurofins Genomics (INVIEW Transcriptome Bacteria) for quality control, rRNA depletion (NEBNext® rRNA Depletion Kit (Bacteria), New England Biolabs), cDNA library preparation and sequencing (Illumina NovaSeq 6000 S4 PE150 XP). Across all samples, an average of 23.7 ± 3.1 million reads were obtained.

### RNA-Seq and differential gene expression analysis

Trimming, mapping and read counting was done in QIAGEN CLC Genomic Workbench (v22) using the “RNA Seq-Analysis” tool run in default settings (length and similarity fraction = 0.8) to obtain gene expression table files with mapped counts. Annotated reference gnomes were made by annotating genome FASTA files using GFF3 files retrieved from Anvi’o contigs databases using the “anvi-get-sequences-for-gene-calls” command with “--export-gff3” flag. Trimmed reads were mapped to annotated reference genomes. When two reads are equally likely to be mapped to two or more positions, the CLC pipeline randomly maps the read to one of the candidate positions and labels the mapped read as “non-unique”. An average of 10.7 ± 1.3 million reads were mapped per sample, of which 9.9 ± 1.6 million reads (93 %) were uniquely mapped. In R (v4.3.1), pangenome gene cluster metadata was mapped to gene expression table by matching Prodigal gene call IDs (see function “r2_RNA_merge_pan_and_count”). Differential gene expression analysis was performed using DESeq2 (v1.40.2) with default settings using pooled counts with a *P*-value threshold of 0.05 and Benjamini and Hochberg adjustment (default to DESeq2) to correct for multiple testing. To allow cross-species comparison of DEG profiles, counts mapped to gene calls within the same gene cluster was pooled for each species.

### Statistical analysis

Principal Component Analysis and Procrustes Superimposition analysis was done using the Vegan (v.2.6-4) package in R. The m^2^ statistic from PS analysis (scale and symmetric true) was tested for significance by a permutation approach (n = 719, maximum number of iterations). Briefly, observations in one matrix are randomly reordered while maintaining the covariance structure within the matrix and a test statistic is calculated and recorded enough times to obtain a sizeable null distribution. A *P-*value for each statistic is then calculated, representing the probability of obtaining a statistic with a value equal to or more extreme of the experimental value.

## DATA AND CODE AVAILABILITY

Experimental data files and R MarkDown scripts used for analysis and plotting are publicly available online on the Open Science Framework database as part of the project name *Chan.Stutz.Pangenome.Chassis* (https://osf.io/yx43n/). Genome and bacterial strain accession numbers can be found in Supplementary Materials. Accession for RNA sequencing data will be made available in the European Nucleotide Archive once ready.

## ACKNOWLEDGEMENTS

We thank Geoff S. Baldwin from the Imperial College of London for their donation of BASIC parts. We thank the SEVA repository for their donation of pSEVA231 plasmid.

## AUTHOR CONTRIBUTIONS

D.T.C.C designed and conducted experiments and wrote the manuscript. H.B designed experiments and wrote the manuscript.

