## Supplementary Figures S1 - S4 for "Pangenomic Landscapes Shape Performances of a Synthetic Genetic Circuit Across *Stutzerimonas* Species"

**Supplementary Figure S1 - S4**

Dennis Tin Chat Chan[1], Hans C. Bernstein[1,2]*

[1]Faculty of Biosciences, Fisheries and Economics, UiT - The Arctic University of Norway, 9019, Tromsø, Norway

[2]The Arctic Centre for Sustainable Energy, UiT - The Arctic University of Norway, 9019, Tromsø, Norway

**Keywords:**

Synthetic Biology, Genetic Inverter, Transcriptomics, Chassis-Effect, Biodesign, Procrustean Superimposition, Context Dependence, Non-Model Organism

### Supplementary Figure S1

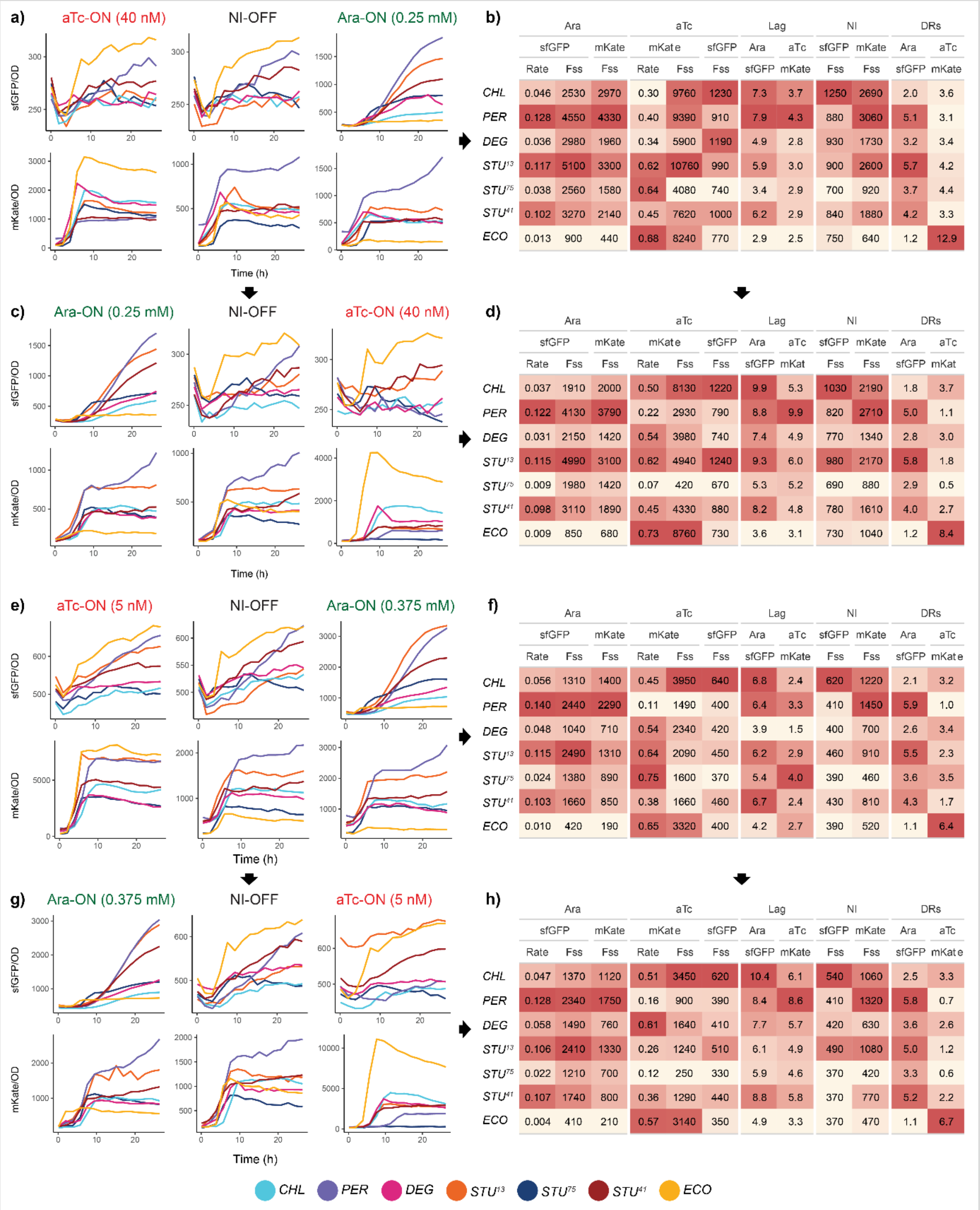


b)

| | Ara | | | aTc | | | Lag | | NI | | DRs | |
|---|---|---|---|---|---|---|---|---|---|---|---|---|
| | sfGFP | | mKate | mKate | | sfGFP | Ara | aTc | sfGFP | mKate | Ara | aTc |
| | Rate | Fss | Fss | Rate | Fss | Fss | sfGFP | mKate | Fss | Fss | sfGFP | mKate |
| *CHL* | 0.046 | 2530 | 2970 | 0.30 | 9760 | 1230 | 7.3 | 3.7 | 1250 | 2690 | 2.0 | 3.6 |
| *PER* | 0.128 | 4550 | 4330 | 0.40 | 9390 | 910 | 7.9 | 4.3 | 880 | 3060 | 5.1 | 3.1 |
| *DEG* | 0.036 | 2980 | 1960 | 0.34 | 5900 | 1190 | 4.9 | 2.8 | 930 | 1730 | 3.2 | 3.4 |
| $STU^{13}$ | 0.117 | 5100 | 3300 | 0.62 | 10760 | 990 | 5.9 | 3.0 | 900 | 2600 | 5.7 | 4.2 |
| $STU^{75}$ | 0.038 | 2560 | 1580 | 0.64 | 4080 | 740 | 3.4 | 2.9 | 700 | 920 | 3.7 | 4.4 |
| $STU^{41}$ | 0.102 | 3270 | 2140 | 0.45 | 7620 | 1000 | 6.2 | 2.9 | 840 | 1880 | 4.2 | 3.3 |
| *ECO* | 0.013 | 900 | 440 | 0.68 | 8240 | 770 | 2.9 | 2.5 | 750 | 640 | 1.2 | 12.9 |

d)

| | Ara | | | aTc | | | Lag | | NI | | DRs | |
|---|---|---|---|---|---|---|---|---|---|---|---|---|
| | sfGFP | | mKate | mKate | | sfGFP | Ara | aTc | sfGFP | mKate | Ara | aTc |
| | Rate | Fss | Fss | Rate | Fss | Fss | sfGFP | mKat | Fss | Fss | sfGFP | mKat |
| *CHL* | 0.037 | 1910 | 2000 | 0.50 | 8130 | 1220 | 9.9 | 5.3 | 1030 | 2190 | 1.8 | 3.7 |
| *PER* | 0.122 | 4130 | 3790 | 0.22 | 2930 | 790 | 8.8 | 9.9 | 820 | 2710 | 5.0 | 1.1 |
| *DEG* | 0.031 | 2150 | 1420 | 0.54 | 3980 | 740 | 7.4 | 4.9 | 770 | 1340 | 2.8 | 3.0 |
| $STU^{13}$ | 0.115 | 4990 | 3100 | 0.62 | 4940 | 1240 | 9.3 | 6.0 | 980 | 2170 | 5.8 | 1.8 |
| $STU^{75}$ | 0.009 | 1980 | 1420 | 0.07 | 420 | 670 | 5.3 | 5.2 | 690 | 880 | 2.9 | 0.5 |
| $STU^{41}$ | 0.098 | 3110 | 1890 | 0.45 | 4330 | 880 | 8.2 | 4.8 | 780 | 1610 | 4.0 | 2.7 |
| *ECO* | 0.009 | 850 | 680 | 0.73 | 8760 | 730 | 3.6 | 3.1 | 730 | 1040 | 1.2 | 8.4 |

f)

| | Ara | | | aTc | | | Lag | | NI | | DRs | |
|---|---|---|---|---|---|---|---|---|---|---|---|---|
| | sfGFP | | mKate | mKate | | sfGFP | Ara | aTc | sfGFP | mKate | Ara | aTc |
| | Rate | Fss | Fss | Rate | Fss | Fss | sfGFP | mKate | Fss | Fss | sfGFP | mKate |
| *CHL* | 0.056 | 1310 | 1400 | 0.45 | 3950 | 640 | 6.8 | 2.4 | 620 | 1220 | 2.1 | 3.2 |
| *PER* | 0.140 | 2440 | 2290 | 0.11 | 1490 | 400 | 6.4 | 3.3 | 410 | 1450 | 5.9 | 1.0 |
| *DEG* | 0.048 | 1040 | 710 | 0.54 | 2340 | 420 | 3.9 | 1.5 | 400 | 700 | 2.6 | 3.4 |
| $STU^{13}$ | 0.115 | 2490 | 1310 | 0.64 | 2090 | 450 | 6.2 | 2.9 | 460 | 910 | 5.5 | 2.3 |
| $STU^{75}$ | 0.024 | 1380 | 890 | 0.75 | 1600 | 370 | 5.4 | 4.0 | 390 | 460 | 3.6 | 3.5 |
| $STU^{41}$ | 0.103 | 1660 | 850 | 0.38 | 1660 | 460 | 6.7 | 2.4 | 430 | 810 | 4.3 | 1.7 |
| *ECO* | 0.010 | 420 | 190 | 0.65 | 3320 | 400 | 4.2 | 2.7 | 390 | 520 | 1.1 | 6.4 |

h)

| | Ara | | | aTc | | | Lag | | NI | | DRs | |
|---|---|---|---|---|---|---|---|---|---|---|---|---|
| | sfGFP | | mKate | mKate | | sfGFP | Ara | aTc | sfGFP | mKate | Ara | aTc |
| | Rate | Fss | Fss | Rate | Fss | Fss | sfGFP | mKate | Fss | Fss | sfGFP | mKate |
| *CHL* | 0.047 | 1370 | 1120 | 0.51 | 3450 | 620 | 10.4 | 6.1 | 540 | 1060 | 2.5 | 3.3 |
| *PER* | 0.128 | 2340 | 1750 | 0.16 | 900 | 390 | 8.4 | 8.6 | 410 | 1320 | 5.8 | 0.7 |
| *DEG* | 0.058 | 1490 | 760 | 0.61 | 1640 | 410 | 7.7 | 5.7 | 420 | 630 | 3.6 | 2.6 |
| $STU^{13}$ | 0.106 | 2410 | 1330 | 0.26 | 1240 | 510 | 6.1 | 4.9 | 490 | 1080 | 5.0 | 1.2 |
| $STU^{75}$ | 0.022 | 1210 | 700 | 0.12 | 250 | 330 | 5.9 | 4.6 | 370 | 420 | 3.3 | 0.6 |
| $STU^{41}$ | 0.107 | 1740 | 800 | 0.36 | 1290 | 440 | 8.8 | 5.8 | 370 | 770 | 5.2 | 2.2 |
| *ECO* | 0.004 | 410 | 210 | 0.57 | 3140 | 350 | 4.9 | 3.3 | 370 | 470 | 1.1 | 6.7 |

**Supplementary Figure S1. Certain hosts experience consistent mKate output attenuation across differing induction schemes.** a) Normalized fluorescence dynamics of toggle assay with induction scheme of 0.25 mM Ara and 40 nM aTc. Initial OFF cells were diluted to respective induction states. Hosts are color coded. (Right) Estimated toggle assay metrics for each host. Color scale is relative to each column. b) (Left) Normalized fluorescence curves of toggled cells diluted to respective opposite inducer. (Right) Estimated toggle assay metrics for each host. Color scale is relative to each column. c) (Left) Normalized fluorescence dynamics of toggle assay with induction scheme of 0.375 mM Ara and 5 nM aTc, note differences in scale of y-axis. Initial OFF cells were diluted to respective induction states. Hosts are color coded. (Right) Estimated toggle assay metrics for each host. Color scale is relative to each column. d) (Left) Normalized fluorescence curves of toggled cells diluted to respective opposite inducer. (Right) Estimated toggle assay metrics for each host. Color scale is relative to each column. NI = No induction, Fss = Late phase steady-state fluorescence, Rate = max specific growth rate. DR = Dynamic Range.

**Supplementary Figure** S2

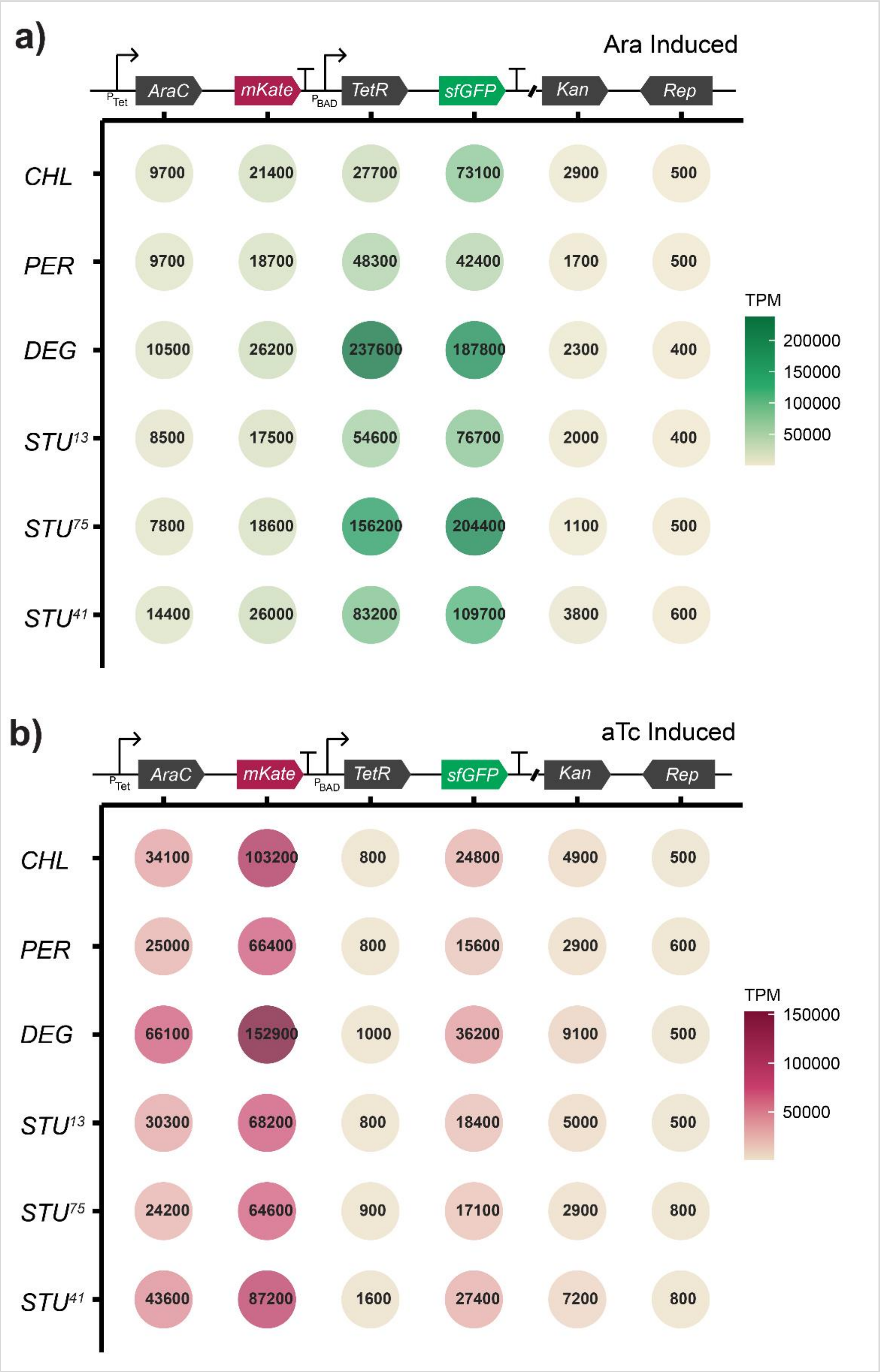


**Supplementary Figure S2. Proportion of mapped transcript abundance to genes encoded in genetic inverter verifies device function.** Relative proportion of mapped transcript abundances, measured as Transcripts per Million (TPM), for Ara and aTc induced host cells. TPM, unlike RPKM, standardizes the denominator used to scale the proportion of mapped reads to a gene, allowing comparison of TPM values within a sample (host).

#### Supplementary Figure S3

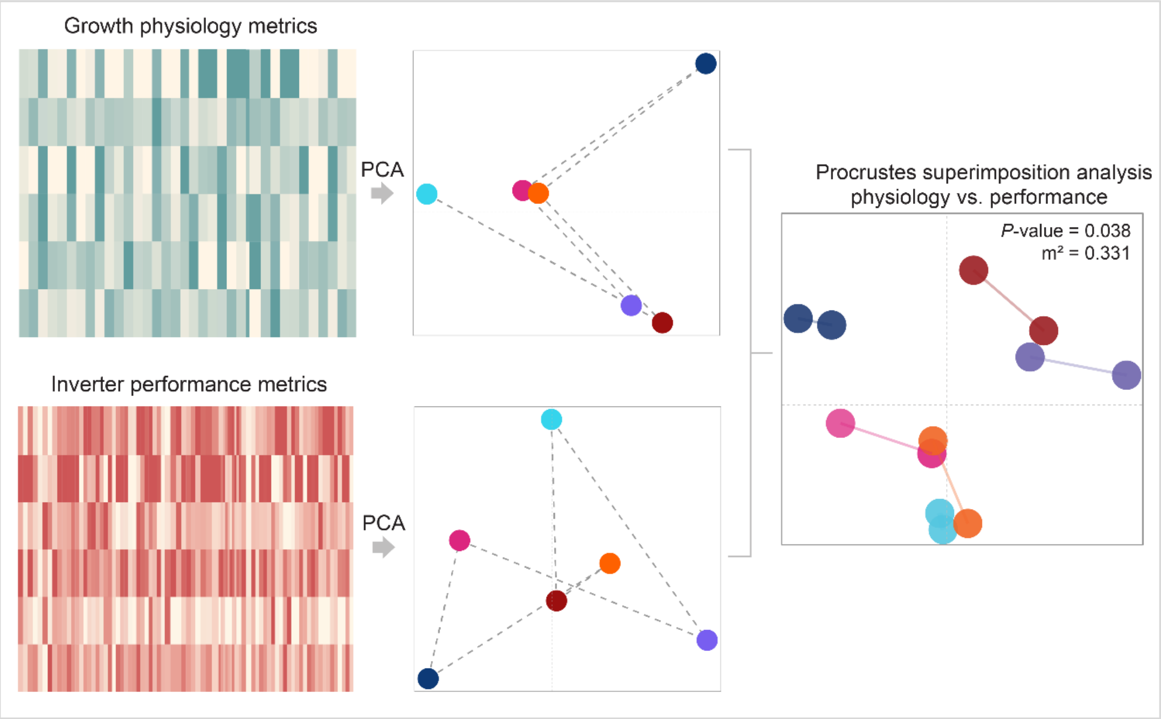


**Supplementary Figure S3. Similarities in growth physiology metrics significantly correlates with similarities in inverter performance metrics within closely related *Stutzerimonas* hosts.** Schematic illustration of PS analysis comparing host similarity in terms of physiological growth metrics and inverter performance metrics.

### Supplementary Figure S4

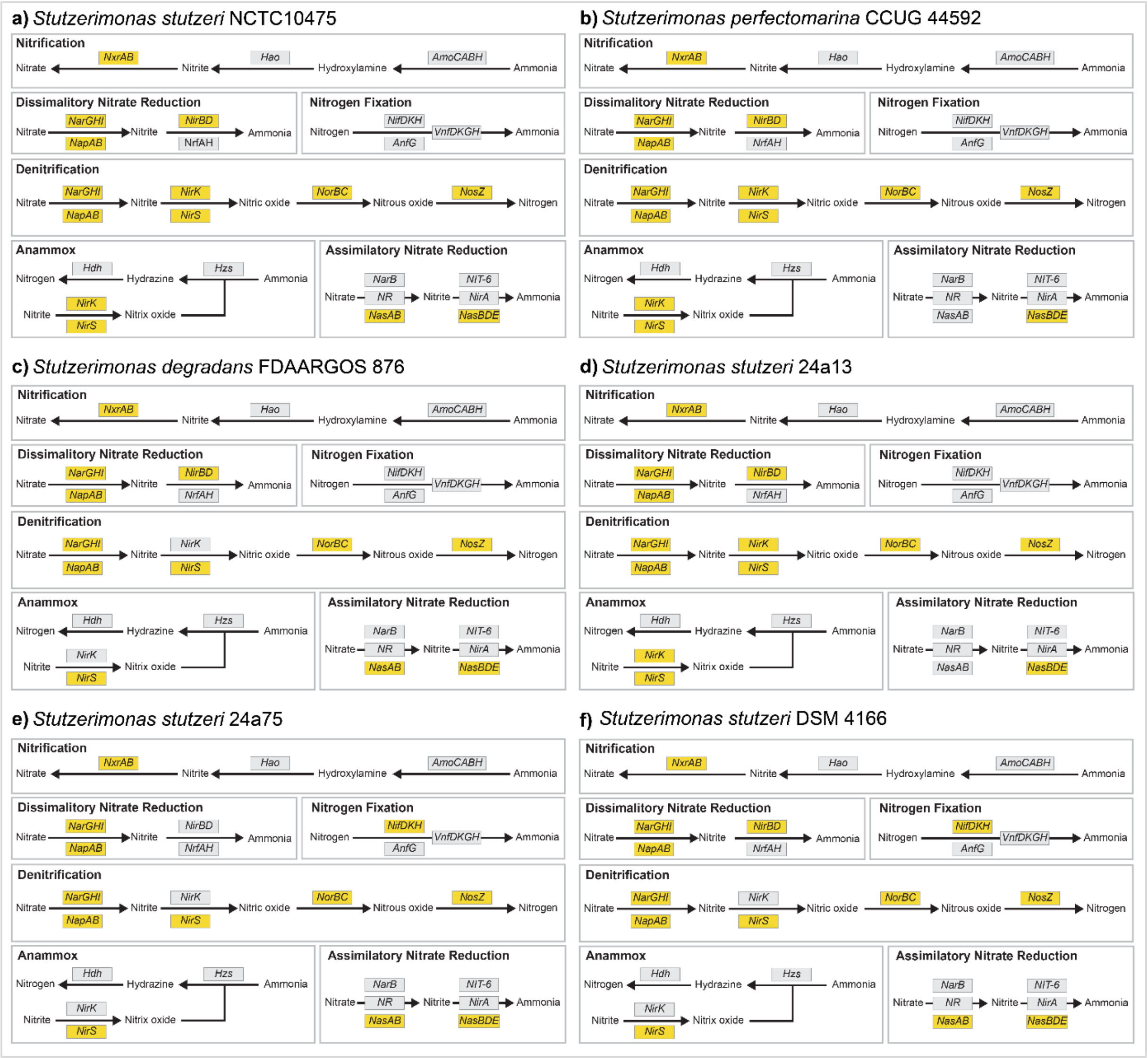


**Supplementary Figure S4. Genes involved in denitrification pathway are inferred to be present in genome of all *Stutzerimonas* hosts while no genes involved in nitrification pathway is found.** a) – f) Overview of presence and absence of genes involved in nitrogen metabolism across *Stutzerimonas* hosts as inferred by KEGG Metabolic Pathway Mapper (https://www.genome.jp/kegg/mapper/) using gene calls provided by Prodigal. Filled in yellow box indicates inferred presence of gene encoding enzyme catalyzing the reaction step and empty (grey) box indicates absence.
